# Using structural analysis *in silico* to assess the impact of missense variants in MEN1

**DOI:** 10.1101/661512

**Authors:** Richard C. Caswell, Martina M. Owens, Adam C. Gunning, Sian Ellard, Caroline F. Wright

## Abstract

Despite the rapid expansion in recent years of databases reporting either benign or pathogenic genetic variation, the interpretation of novel missense variants can remain challenging, particularly for clinical or genetic testing laboratories where functional analysis is often unfeasible. Previous studies have shown that thermodynamic analysis of protein structure *in silico* can discriminate between groups of benign and pathogenic missense variants. However, although structures exist for many human disease-associated proteins, such analysis remains largely unexploited in clinical laboratories. Here, we analysed the predicted effect of 338 known missense variants on the structure of Menin, the *MEN1* gene product. Results provided strong discrimination between pathogenic and benign variants, with a threshold of >4 kcal/mol for the predicted change in stability providing a strong indicator of pathogenicity. Subsequent analysis of 7 novel missense variants identified during clinical testing of MEN1 patients showed that all 7 were predicted to destabilise Menin by >4 kcal/mol. We conclude that structural analysis provides a useful tool in understanding the impact of missense variants in *MEN1*, and that integration of proteomic with genomic data could potentially contribute to the classification of novel variants in this disease.

## INTRODUCTION

The rapid expansion in recent years of genomic data from both patient and control groups has vastly improved the quantity and quality of information that is available to clinicians in attempting to classify novel genetic variants. While it is often straightforward to interpret likely loss-of-function variants such as stop-gain or frameshift variants, the same is not true of missense variants, where the effect of an amino acid substitution is likely to be specific to its context in the protein of interest. Moreover, such variants are often rare or unique, and thus must be interpreted on a case-by-case basis.

Numerous methods have been developed for predicting the phenotypic effect of missense variants. As has been comprehensively reviewed elsewhere (1), these methods rely either on analysis of DNA and protein conservation, protein structure-based analysis, or a combination of the two. In the case of the latter, widely used tools such as PolyPhen are able to incorporate information on the nature of the amino acid change itself (e.g. Grantham distance between native and variant amino acids, changes in polarity or charge), effects on predicted secondary structure and, where available, data derived from the structural context, such as changes in hydrogen bonding or atomic crowding.

However, such data is used in a qualitative, rule-based manner in the final prediction (1), and the tools which are most widely used in the clinical setting do not specifically address the quantitative effects of missense variants on protein stability. Nevertheless, these effects can be calculated where there is an experimental or modeled 3D structure for the protein of interest, and programs such as FoldX (2), Rosetta (3, 4) or other computational methods have been widely used by structural biologists to investigate the effects of missense variants on protein folding and stability (5, 6).

Despite this, few studies have sought to address whether there is a direct clinical application of such an approach, i.e. whether pathogenic and benign variants can be distinguished on the basis of their predicted effects on thermodynamic stability.

The potential utility of protein stability data towards the analysis of missense variants has recently been demonstrated in studies of the Lynch syndrome protein, MSH2 (7), and in phenylalanine hydroxylase (PAH) (8), in which pathogenic variants result in phenylketonuria. Both these studies combined *in silico* analysis with extensive functional analysis of a number of *MSH2* and *PAH* variants; however, resources for the latter are unlikely to be routinely available in clinical genetics laboratories. We have therefore asked whether *in silico* analysis, based predominantly on the predicted effects of missense variants on protein stability, can help discriminate between pathogenic and benign variation in the context of clinical testing of the *MEN1* gene.

Pathogenic variants in the *MEN1* gene cause Multiple Endocrine Neoplasia type I, an autosomal dominant disorder, in which patients develop neoplastic lesions in various endocrine tissues, principally the parathyroids, pituitary and pancreas (9). Pathogenic variants may either be inherited or acquired, but in both cases development of disease requires loss of heterozygosity consistent with a role for the *MEN1* gene product as a tumor suppressor. The most common presenting feature of MEN1 is hyperparathyroidism, which occurs in ∼95% of patients due to tumors of the parathyroid gland; however, tumors are also frequently observed in the pancreatic islets (40-70%) and pituitary (30-40%) (10). Patients may also develop tumors of the adrenal cortex, carcinoid tumors and non-endocrine tumors, including lipomas, angiofibromas, collagenomas and meningiomas (11), resulting in a range of clinical symptoms which may overlap with other diseases of different genetic etiology (12–14). This overlap presents one of the key problems in assessing genetic variants in cases of MEN1. While a large number of pathogenic variants in *MEN1* have been reported, genetic testing continues to uncover novel missense substitutions which require assessment of their potential pathogenicity. A further confounding issue is the often later onset of disease, with reported age-related penetrance of 10-43% at 20 years and 81-94% by 50 years (10, 15), which may lead to apparent non-segregation of a variant with disease within a family pedigree.

The identification of a genetic etiology has important implications for the patient and for their family members. With the exception of pituitary neuroendocrine tumors, MEN1-associated tumors are usually multiple and treatment is therefore challenging, requiring a multi-disciplinary team of experts to reduce morbidity and mortality (16). The identification of the familial disease-causing variant enables the identification of carriers when they are still asymptomatic. Clinical surveillance in these individuals allows early recognition of the clinical manifestations and therapeutic intervention. For example, primary hyperparathyroidism often remains asymptomatic in many patients but prolonged hypercalcemia usually results in bone loss and/or nephrocalcinosis (17).

Approximately 20% of the variants identified in the *MEN1* gene are missense variants (18). The standards and guidelines published by the American College of Medical Genetics and Genomics (ACMG) and the Association for Molecular Pathology (AMP) describe a framework for the classification of sequence variants (19). Adjustments to this framework for the interpretation of *MEN1* missense variants has been proposed (20). However both agree that variants of uncertain significance should not be used to guide the clinical management of patients. This could lead to an under-diagnosis of MEN1 and a lost opportunity for screening at-risk relatives. For these reasons, methods to assist the classification of variants in *MEN1* would be of clinical value. The availability of a number of experimental structures for Menin, the *MEN1* gene product, raises the possibility that structural analysis may provide such clinical utility.

We report here that thermodynamic analysis of *MEN1* variants *in silico* provides a very strong positive predictive value for pathogenicity, thereby helping to assess the impact of novel missense variants on protein function and potentially allowing its use as an aid to variant classification, and discuss briefly the scope for wider application of this approach to other diseases.

## MATERIALS & METHODS

### Variant groups, transcripts and numbering

Previously-reported missense SNVs in *MEN1* were downloaded from the Human Gene Mutation Database, Professional version (HGMD Pro) (21), the Genome Aggregation Database (gnomAD) (22) and the Sydney Genomics Collaborative Database (SGCD) (23). For the purposes of this analysis, variants were divided into groups as follows: pathogenic: DM (‘disease mutation’) class variants reported in HGMD Pro but not in gnomAD or SGCD (n=162); benign: variants reported in gnomAD or SGCD but not as DM class in HGMD Pro (n=206); uncertain: variants reported as DM in HGMD Pro and present in gnomAD and/or SGCD (n=14). Different nucleotide substitutions resulting in the same coding change were regarded as a single missense substitution. In addition to these previously-reported variants, analysis was performed on seven novel missense variants: H46P; A164P; L175P; A345P; I360F; F364S; and G419D (see Table 1 for details). These variants were identified in our laboratory as part of the NHS (England) Genetic Testing service for rare inherited diseases. The patients tested fulfilled the criteria for a clinical diagnosis of MEN1 (10), presenting with at least two out of the three main MEN1-associated endocrine lesions or one typical MEN1-associated tumour and a first-degree relative with MEN1 or MEN1-associated lesion at a young age. For patients with a family history, the relevant variants (H46P, A164P, I360F and F364S) were all shown to co-segregate with disease in the family.

**Table 1:** Details of 7 novel missense variants in MEN1. All variants refer to *MEN1* transcript NM_130799.2, protein NP_570711.1 (610 amino acid isoform).

| variant # | 1 | 2 | 3 | 4 | 5 | 6 | 7 |
| --- | --- | --- | --- | --- | --- | --- | --- |
| HGVS c. notation | c.137A>C | c.490G>C | c.524T>C | c.1033G>C | c.1078A>T | c.1091T>C | c.1256G>A |
| HGVS p. notation | p.(His46Pro) | p.(Ala164Pro) | p.(Leu175Pro) | p.(Ala345Pro) | p.(Ile360Phe) | p.(Phe364Ser) | p.(Gly419Asp) |
| genomic variant (GRCh37/hg19) | chr11:64577445T>G | chr11:64575527C>G | chr11:64575493A>G | chr11:64573720C>G | chr11:64573214T>A | chr11:64573201A>G | chr11:64572600C>T |
| reported in gnomAD? | no | no | no | no | no | no | no |
| SIFT prediction | Damaging | Damaging | Damaging | Damaging | Damaging | Damaging | Damaging |
| PROVEAN prediction | Deleterious | Deleterious | Deleterious | Deleterious | Deleterious | Deleterious | Deleterious |
| PolyPhen prediction | probably damaging | probably damaging | probably damaging | probably damaging | probably damaging | probably damaging | probably damaging |
| REVEL score | 0.894 | 0.925 | 0.965 | 0.909 | 0.883 | 0.945 | 0.912 |

Menin, the protein product of the *MEN1* gene, occurs in two major isoforms of 615 or 610 amino acids, which arise by use of alternative splice donor sites in exon 1 such that the shorter isoform lacks residues 149-153 of the longer. While gnomAD and SGCD variants are annotated according to the 615-residue isoform encoded by transcripts NM_130803/ENST00000337652, HGMD Pro and structural databases use the 610-residue isoform encoded by NM_130799/ENST00000312049 as default. All numbering in this manuscript refers to the 610-residue form of Menin, and variants from gnomAD and SGCD have been re-annotated accordingly.

### Protein structures

Structures of human Menin were downloaded as PDB files from the worldwide Protein Data Bank (24); a full list of the 29 crystal structures, containing 31 discrete Menin chains, used in this analysis is shown in Table 2. Any non-native amino acids (e.g. affinity purification tags) in these structures were removed from PDB files prior to further analysis.

**Table 2:**
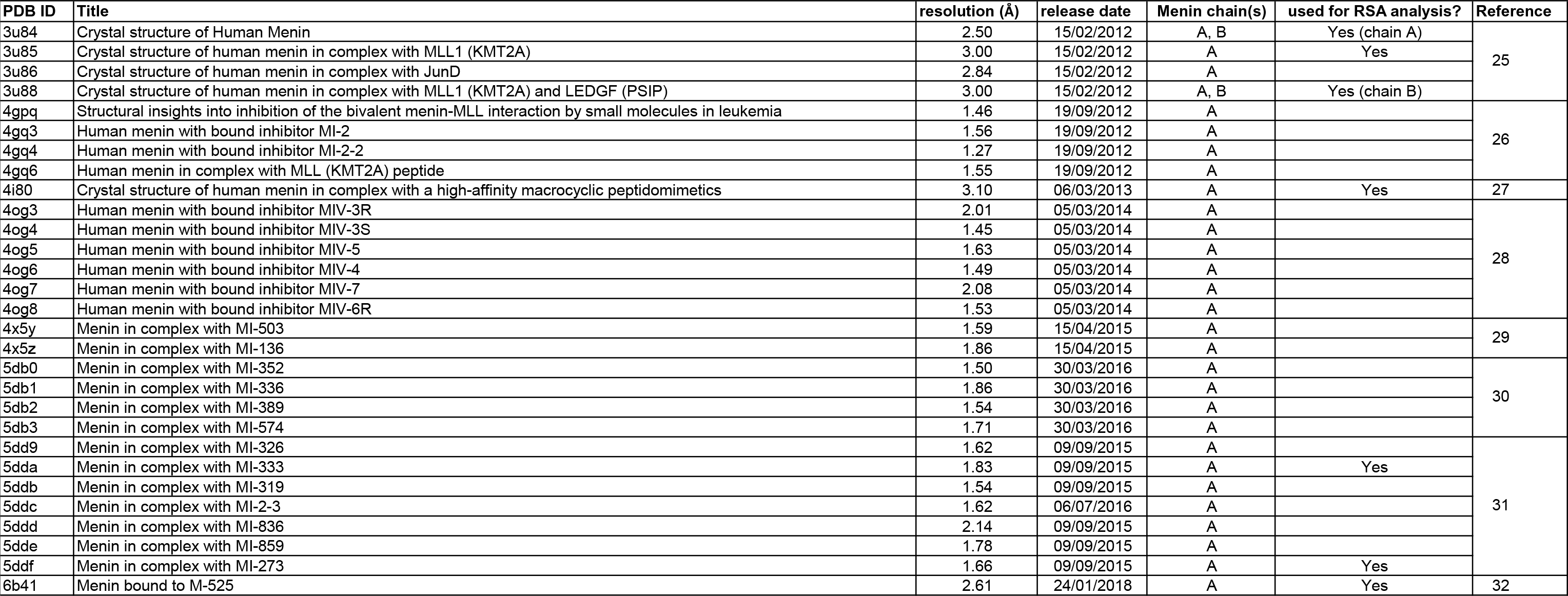
MEN1 crystal structures used in FoldX analysis. A total of 29 PDB structures containing 31 Menin chains were used for thermodynamic analysis using FoldX; 7 representative structures were also used for relative solvent accessibility (RSA) analysis.

### *In silico* mutagenesis and thermodynamic analysis

Prior to *in silico* mutagenesis, sidechain repair and energy minimization was performed on all 31 Menin chains in isolation, using the RepairPDB function of the FoldX modeling suite, version 4 (33). The FoldX BuildModel function was then used to introduce individual substitutions into each of the repaired PDB structures. Of the 389 unique missense variants, 338 were covered by at least one PDB structure (pathogenic, n=161; benign, n=161; uncertain, n=9; novel, n=7). For each substitution, FoldX reported a change in free energy (ΔΔ*G*) resulting from the substitution; from this, an average ΔΔ*G* value was calculated for each variant across all structures containing the relevant position. In total, all 31 structures were used for 308/338 variants (mean for all variants = 29), whereas due to differences in coverage of individual PDB files, analysis was possible using only a single structure for 7 variants. A full list of variants, the number of PDB structures analysed for each and average ΔΔ*G* values for each variant is shown in Table 3. All structures were visualized in PyMOL (34).

**Table 3:**
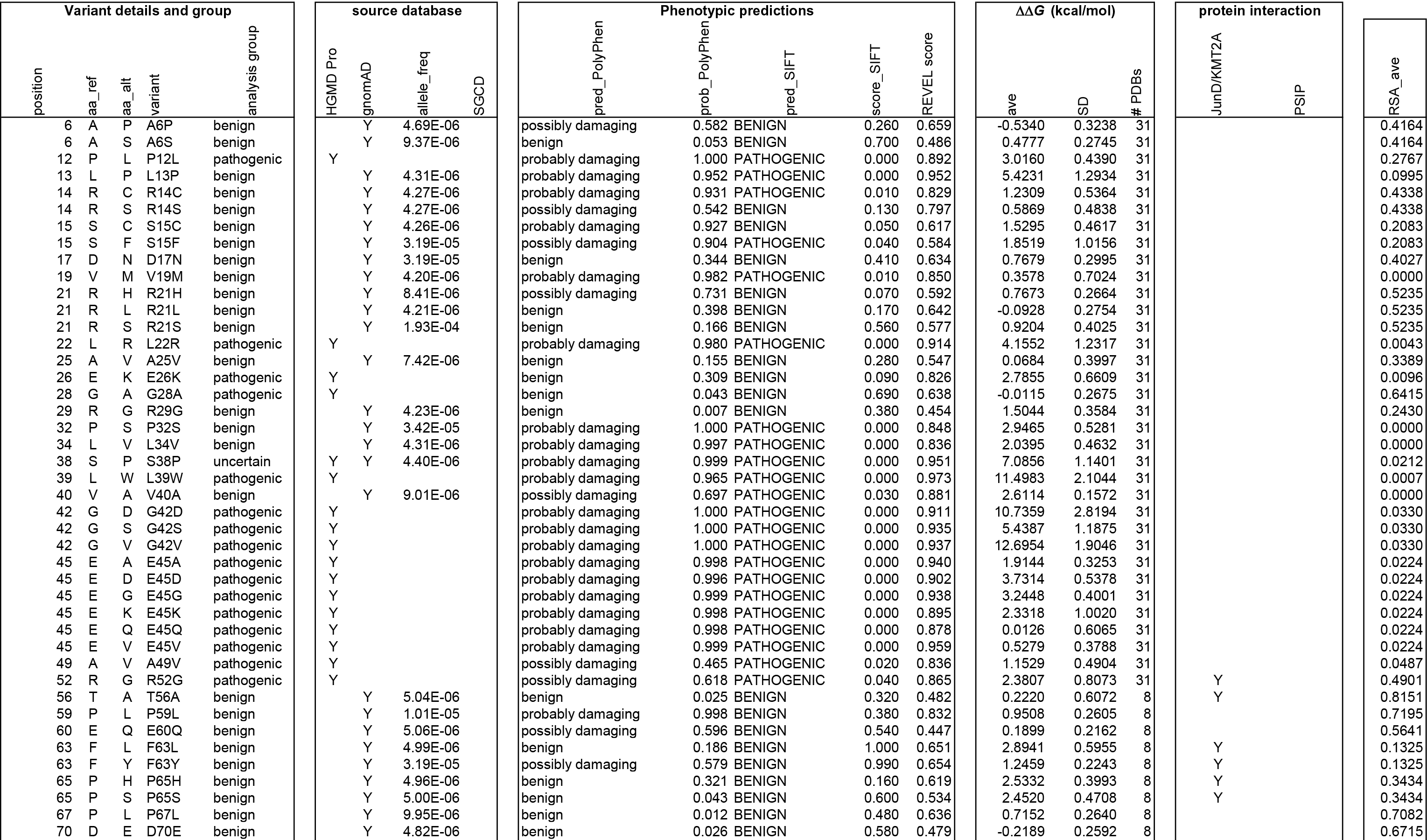
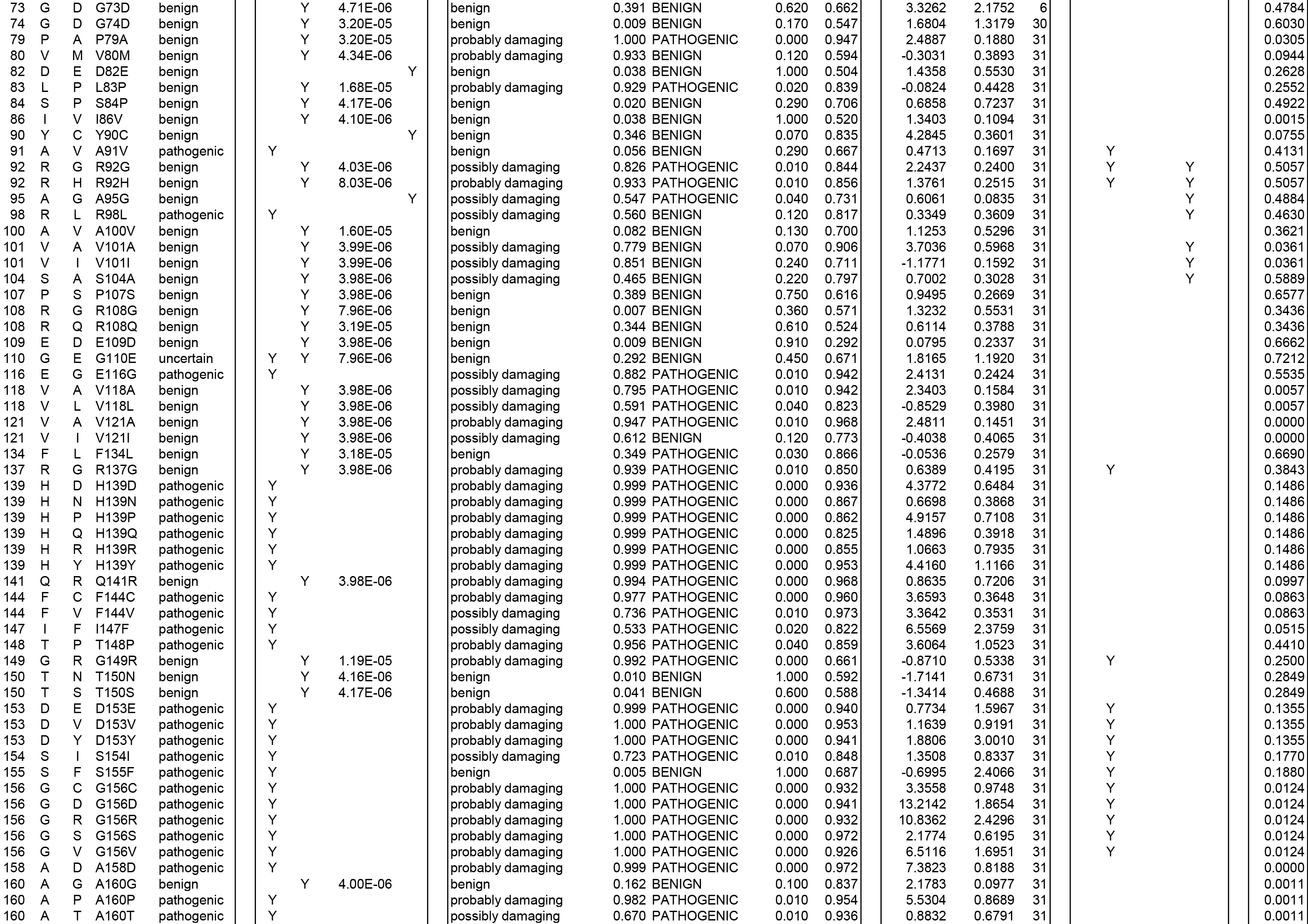
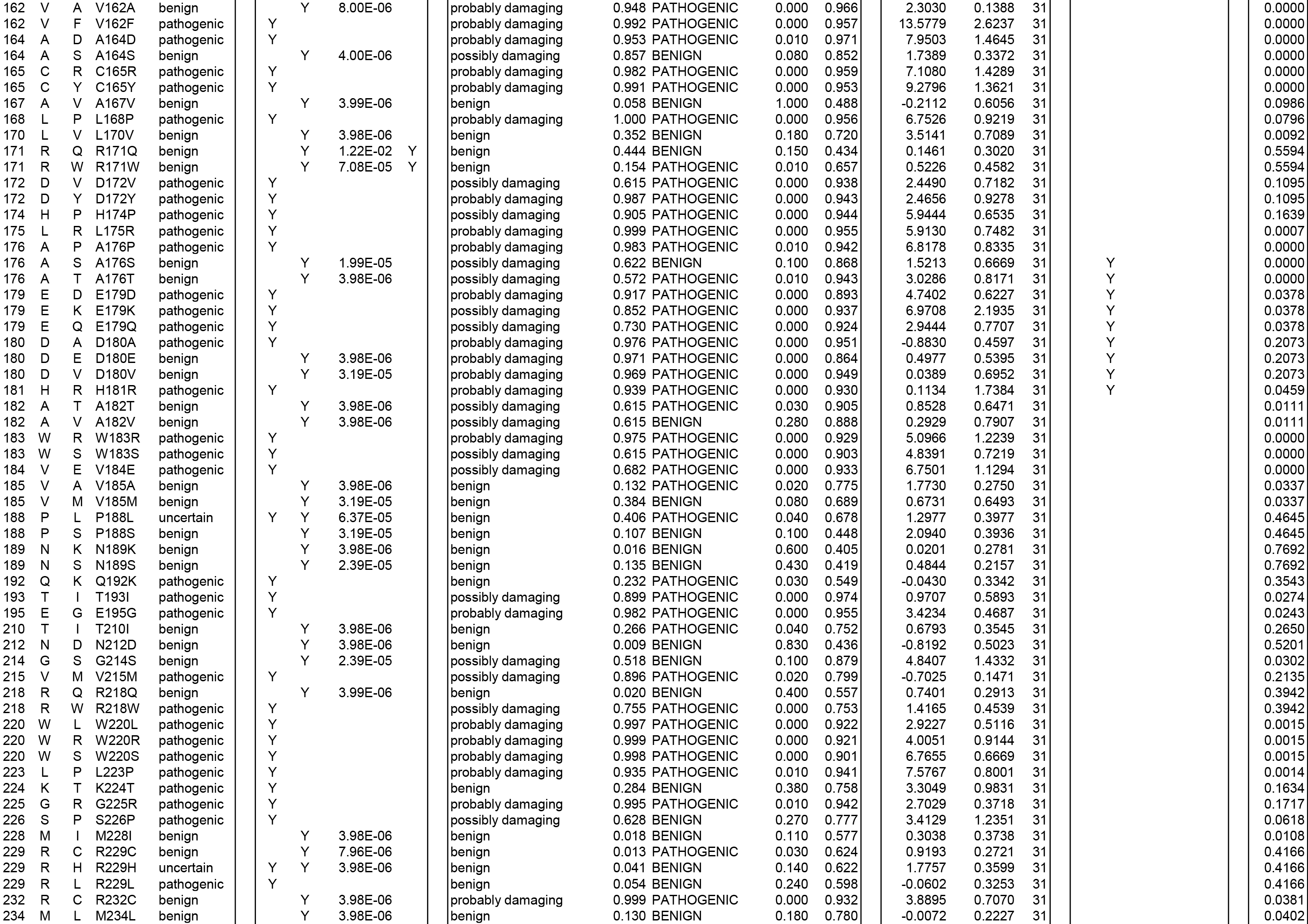
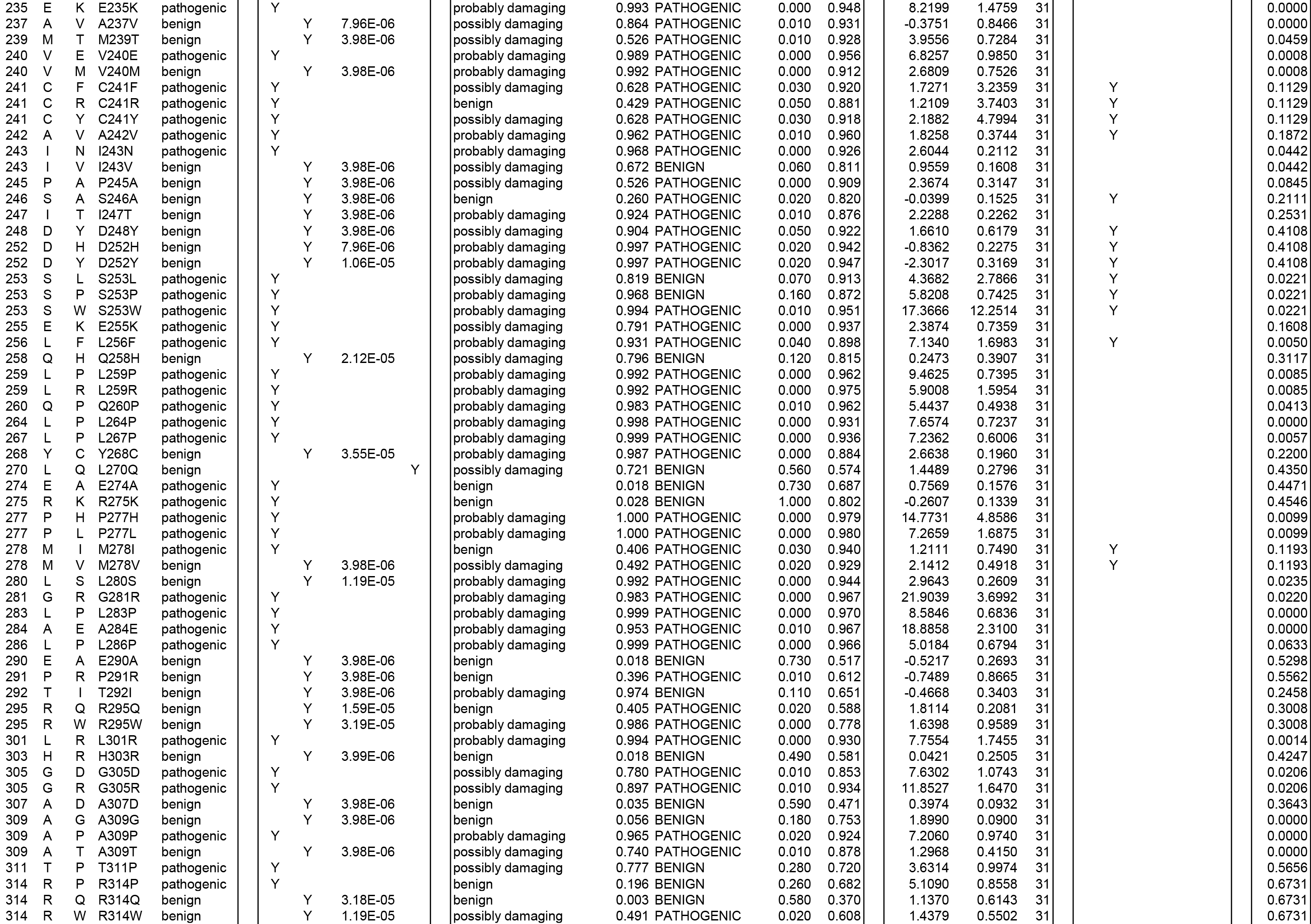
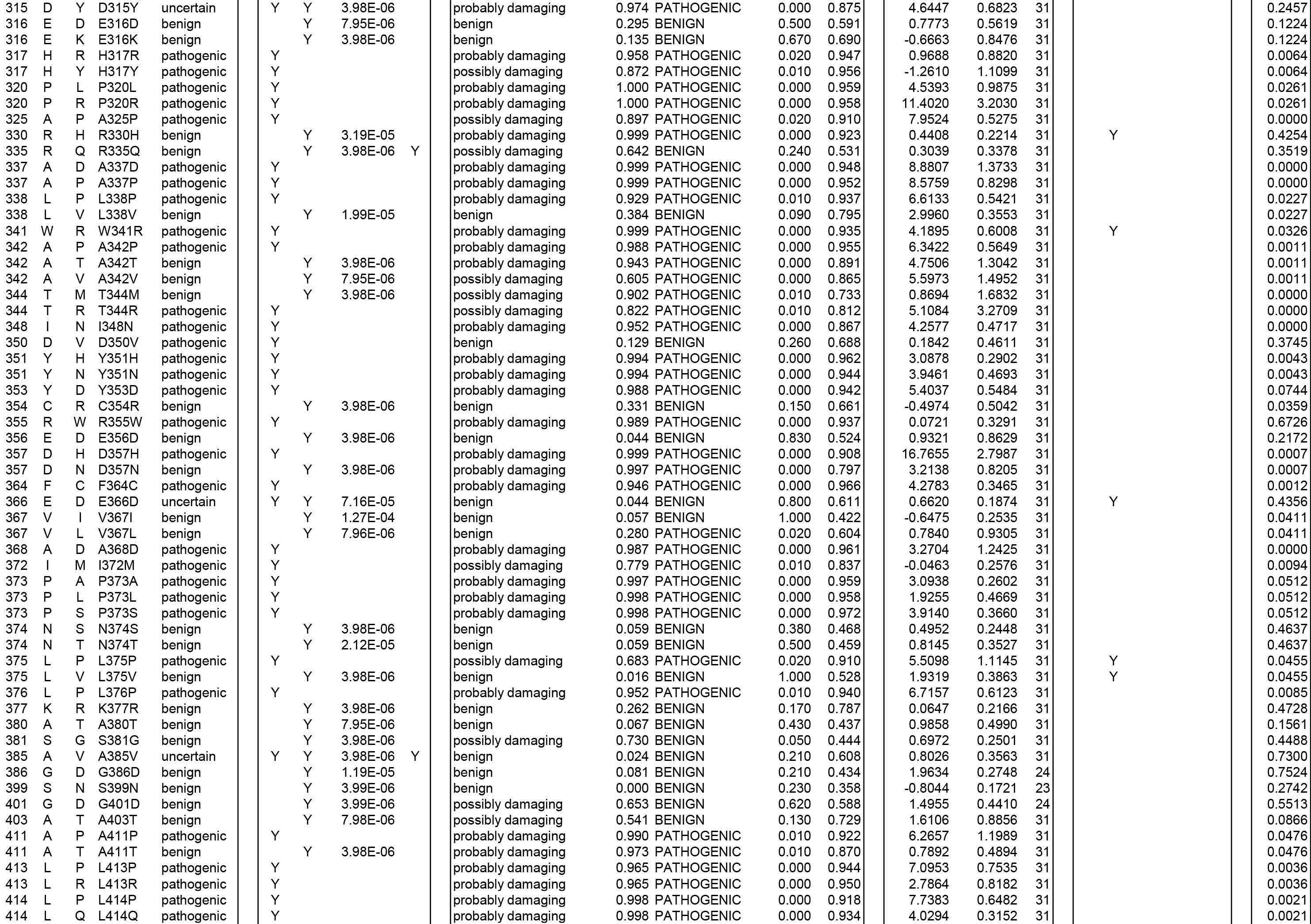
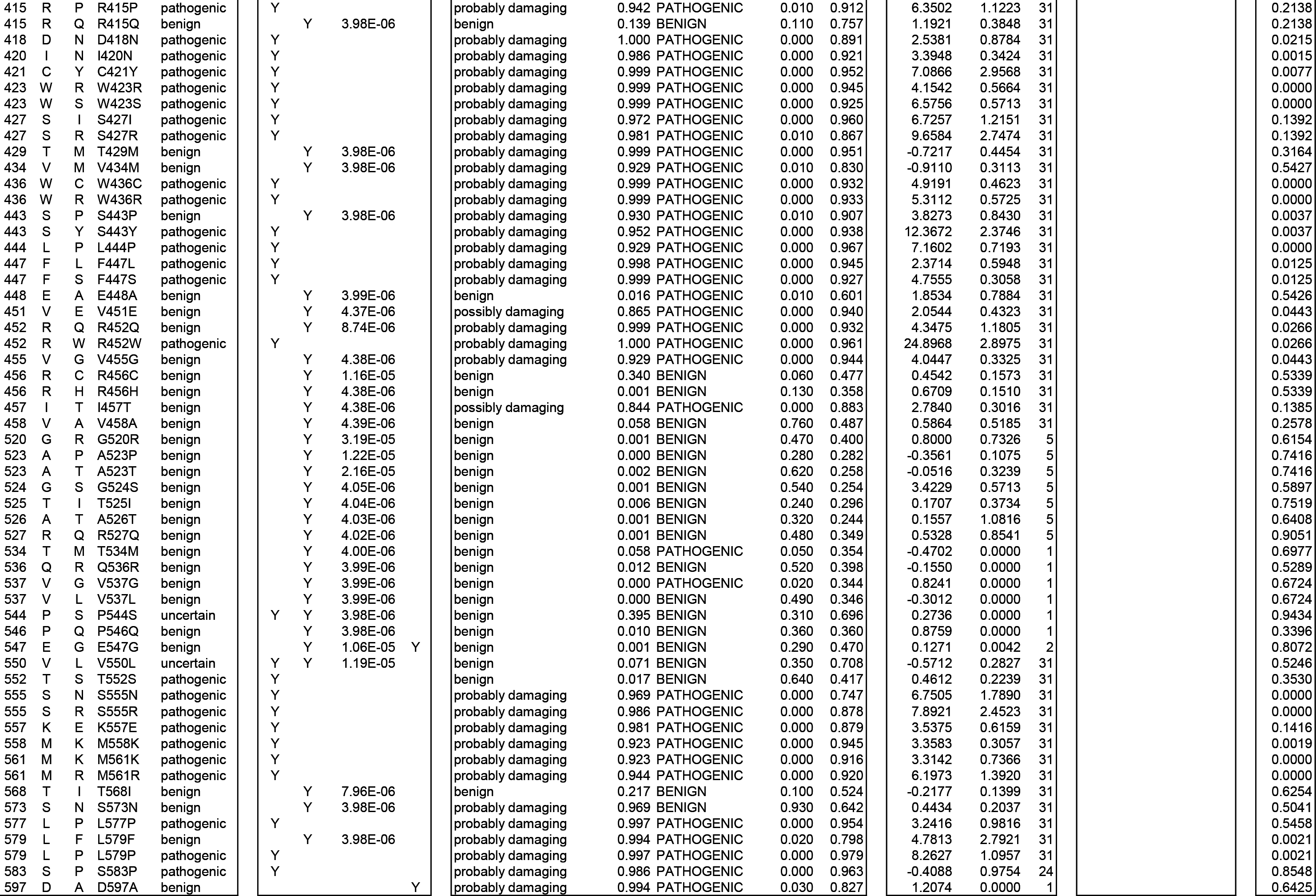
Unique missense variants, FoldX analysis (ΔΔ*G*) and relative solvent accessibility (RSA). Details are shown for unique missense variants in pathogenic (n=161), benign (n=161) and uncertain (n=9) groups as defined in Materials & Methods. Where the source database included gnomAD, frequency is shown for the variant allele. Phenotypic predictions for each variant show prediction and probability data for PolyPhen2, prediction and score for SIFT and score for REVEL prediction. Results of thermodynamic analysis are shown as average ΔΔ*G* value and standard deviation, derived from FoldX calculation using the number of PDB structures indicated (#PDBs). ‘Protein interaction’ columns indicate residues annotated in relevant PDB entries as interacting directly either with JunD and/or KMT2A, both of which bind to the JunD binding pocket of Menin, or to PSIP. Results of relative solvent accessibility (RSA) analysis are shown as average values derived from DSSP analysis of 7 representative structures.

### Calculation of solvent accessibility

The absolute area accessible to solvent (ASA) was calculated on a residue-by-residue basis for 7 representative structures of Menin using DSSP (35, 36) version 3.0.0 (37). After calculating an average ASA value for each residue, relative solvent accessibility (RSA) was derived using the theoretical scale described by Tien et al. (38). A list of structures used for DSSP analysis is included in Table 2.

## RESULTS

### Pathogenic variants in MEN1 are predicted to be destabilizing

Over 30 crystal structures have previously been reported for Menin (e.g. Figure 1A); most of these contain the protein in isolation or bound to a small (drug) ligand, while others show Menin in complex with peptides from JunD, KMT2A or PSIP (Figure 1B; Table 2). Although all structures have been derived from expression of full-length (or near full-length) Menin, a number of regions remain unresolved in crystal structures. These regions predominantly lie in the C-terminal of the protein and correspond to stretches of predicted intrinsic disorder (39) in the protein (Figure 1C, D), presumably resulting in high mobility within crystals. Interestingly, while these regions contain a similar distribution of benign variants as that seen in the protein as a whole, pathogenic variants are rare in regions of predicted disorder (Figure 1D); however, we cannot rule out the possibility that the lack of pathogenic variants in disordered regions is due to reporting bias towards variants which lie close to those already known. As a result of this distribution of pathogenic variants almost entirely within ordered regions, the vast majority (161/162) are covered by one or more PDB entries and are thus amenable to structural analysis.

**Figure 1:**
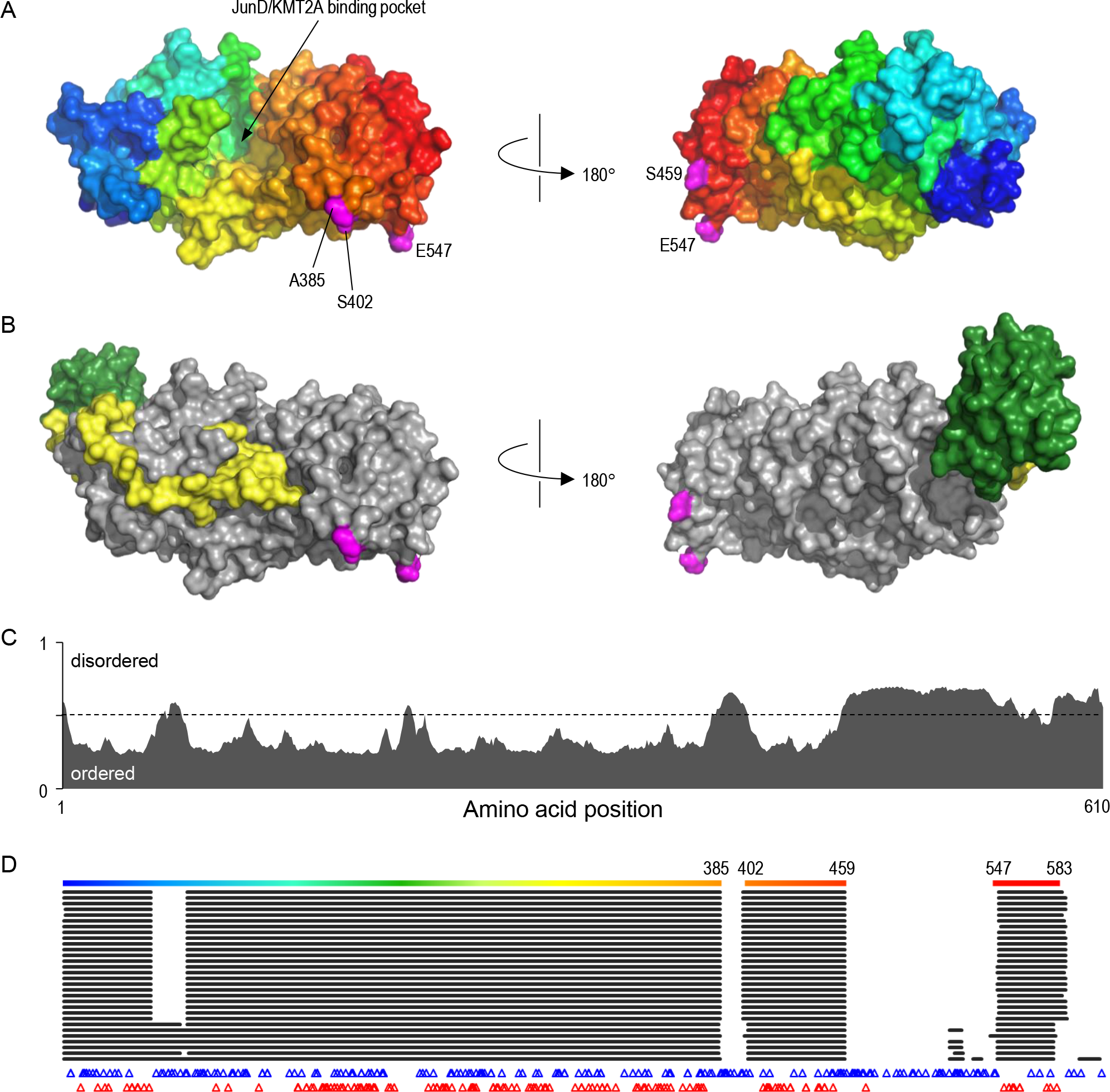
Structure and disorder in Menin. A) The structure of Menin, as represented by PDB entry 3u88 chain A; protein surface is coloured from blue, N-terminal to red, C-terminal; the position of the binding pocket for JunD and KMT2A is indicated; numbered residues coloured magenta indicate positions flanking disordered loops which are not resolved in the crystal structure. B) Menin (grey) in complex with KMT2A (yellow) and PSIP (green), as determined in PDB 3u88; note that while one end of KMT2A occupies the binding pocket, interaction with PSIP and other regions of KMT2A extends over a wider region of the Menin surface. C) Probability of intrinsic disorder in Menin, as calculated by the MetaDisorder predictor, plotted against amino acid position; extended regions of probability >0.5 are considered to be disordered. D) Coverage of Menin residues in the 31 PDB structures used in this analysis, aligned against amino acid position as in part C. The top line shows coverage in PDB 3u88A, coloured as in 1A; numbering indicates residues flanking unstructured regions missing from the crystal structure. Below this, black horizontal lines show coverage for the 30 remaining PDB structures, while positions of benign and pathogenic variants are indicated by blue or red triangles indicate respectively. Note that regions of predicted intrinsic disorder are absent from the majority, if not all crystal structures, consistent with greater mobility of these residues within the crystal, and that few pathogenic variants have been reported in these regions.

The overall structure of Menin is highly comparable within all reported PDB structures (alignment to PDB 6b41 yields an average root-mean-square deviation, RMSD, of 0.65 Å; range 0.55-1.10 Å). Moreover, there is no significant effect of ligand binding on Menin structure (Figure 2). Since different PDB files contain slightly different numbers of amino acids but there are no obvious structural outliers, all available structures were used for thermodynamic analysis of missense variants *in silico* using FoldX.

**Figure 2:**
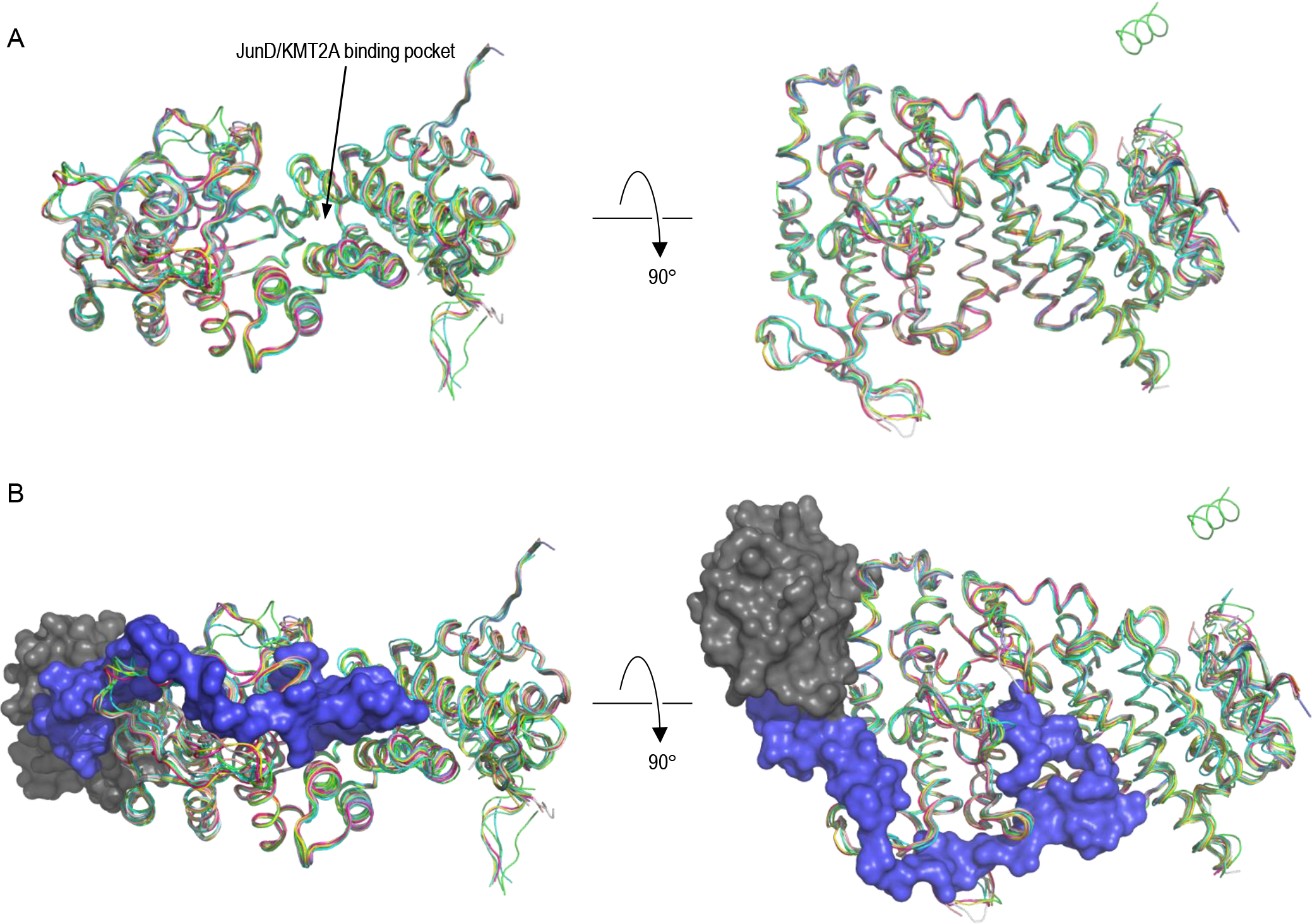
Alignment of Menin structures. A) The α carbon atoms of the 31 Menin structures used in this study were aligned to that of PDB 6b41; each chain is shown in ribbon format, coloured by PDB and chain identifier; the position of the JunD/KMT2A binding pocket is indicated; the short helix visible at the top right of the rotated figure corresponds to residues 596-608 at the extreme C-terminal of Menin, which were resolved only in PDB 3u84 chain A. B) As A, but superimposed with the structures of MLL (blue) and PSIP (grey) from PDB 3u88.

Variant groups were highly distinguishable by their predicted effect on thermodynamic stability, as represented by average ΔΔ*G* value calculated across all structures, with most putatively benign (gnomAD and SGCD) variants having little or no effect (average ΔΔ*G* for all variants, 1.13 kcal/mol; SD, 1.46 kcal/mol), whereas pathogenic (HGMD only) variants were predicted to be strongly destabilizing (average ΔΔ*G*, 5.06 kcal/mol; SD 4.25 kcal/mol) (Figure 3A). Notably, the seven novel missense variants were also predicted to be strongly destabilising (average ΔΔ*G*, 7.67 kcal/mol; SD 3.14 kcal/mol). Analysis of ΔΔ*G* values for individual PDB structures showed a similar separation of putative benign and pathogenic variant groups, with the vast majority of variants falling into a similar range for all structures (Figure 3B). We further compared the effect at multi-allelic sites where different benign and pathogenic missense variants occur at the same position. Analysis of 27 benign and 23 pathogenic variants co-occurring at 22 residues again showed that the difference between the two groups was highly significant (*p* = 0.0002), and that pathogenic missense changes were more strongly destabilizing than benign ones at the same position (average ΔΔ*G* value by group = 6.81 kcal/mol and 2.18 kcal/mol respectively) (Figure 3C).

**Figure 3:**
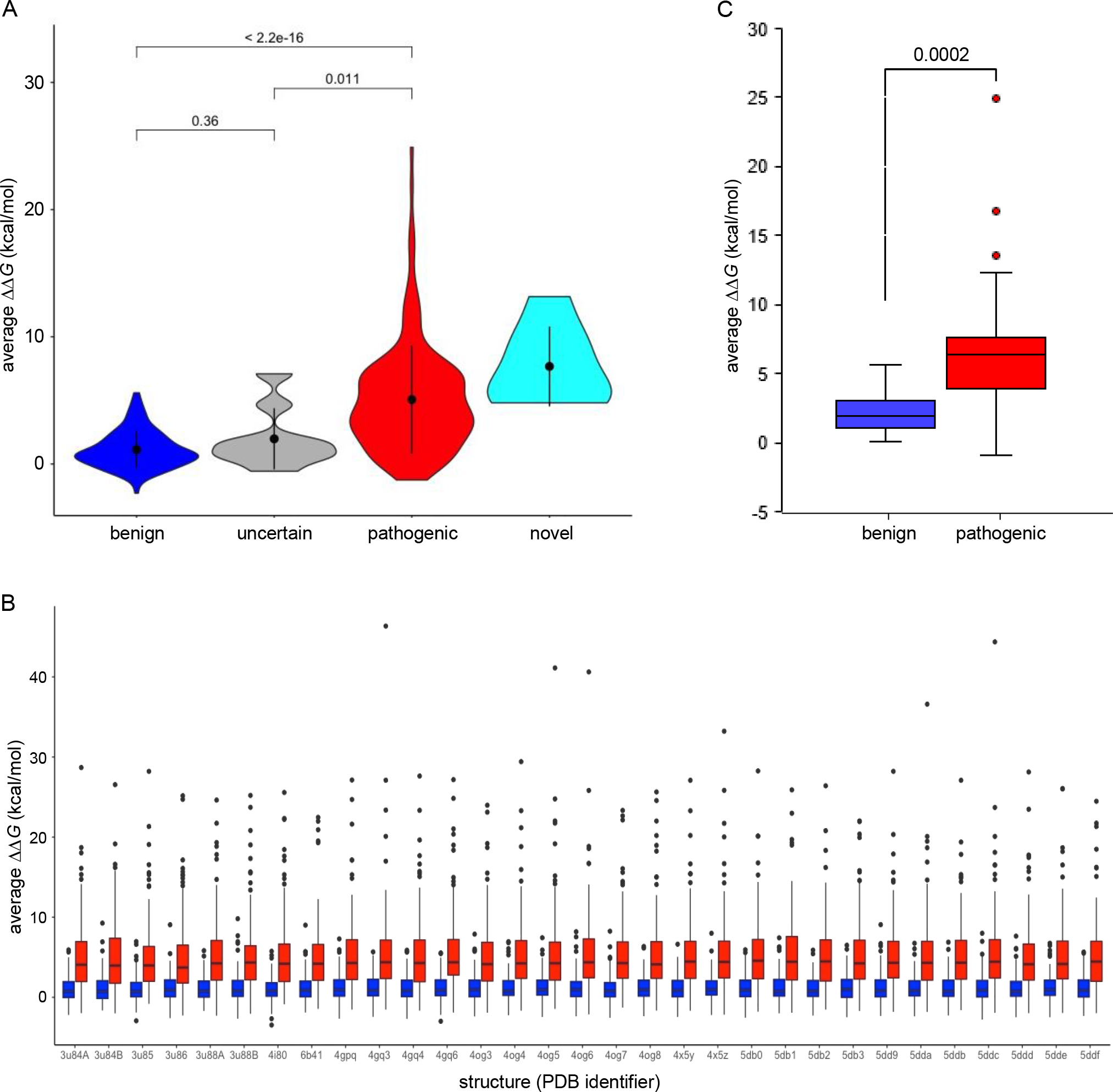
Pathogenic variants are predicted to destabilise Menin structure. A) *In silico* mutagenesis and thermodynamic analysis for Menin variants; for each variant, the average change in thermodynamic stability, ΔΔ*G*, was calculated across all structures contained the relevant residue, then plotted by variant group; black circles and vertical lines within each data area represent median and upper and lower quartiles respectively. Numbering above data points shows *p* values (Student’s Two-tailed t-test) between groups as indicated. B) ΔΔ*G* values for benign (blue) and pathogenic (red) variant groups calculated for 31 individual PDB structures as shown on the x-axis. C) Average ΔΔ*G* values for benign and pathogenic variants occurring at the same amino acid position (residues with one benign and one pathogenic variant, n=16; residues with two benign and one pathogenic variants, n=5; residues with one benign and two pathogenic variants, n=1); coloured boxes show the range between upper and lower quartiles; horizontal lines within each data box show median value; data points are shown for outliers only. The difference in the average ΔΔ*G* value between groups was highly significant (*p*=0.0002).

If variants which destabilize Menin structure do indeed have a greater tendency to be pathogenic, it might be expected that variants most frequently observed in the general population would have the least destabilizing effect. This appears to be the case, as variants with the highest population frequency had average predicted ΔΔ*G* values in the range -1 to +1 (Figure 4); as the error in FoldX calculations is approximately ±0.8 kcal/mol (2), this suggests little or no effect of these variants on protein stability. Notably, those variants which have also been observed in an aging healthy population, as represented by the SGCD cohort (median age, 80-85 years) and are therefore most likely to be truly benign, all occur within this range of ΔΔ*G* values. This group includes the only commonly-occurring missense *MEN1* variant, R171Q, which has an average ΔΔ*G* value of 0.15 kcal/mol. Conversely, we note that some variants reported in gnomAD have ΔΔ*G* values >4 kcal/mol, and in fact 2/9 of these variants (S38P, D315Y) have also been reported as disease-causing in HGMD Pro. This may reflect the confounding effect of late onset of symptoms in MEN1 on apparent constraint against coding variation, whereby some variants reported in gnomAD may in fact lead to disease in later life.

**Figure 4:**
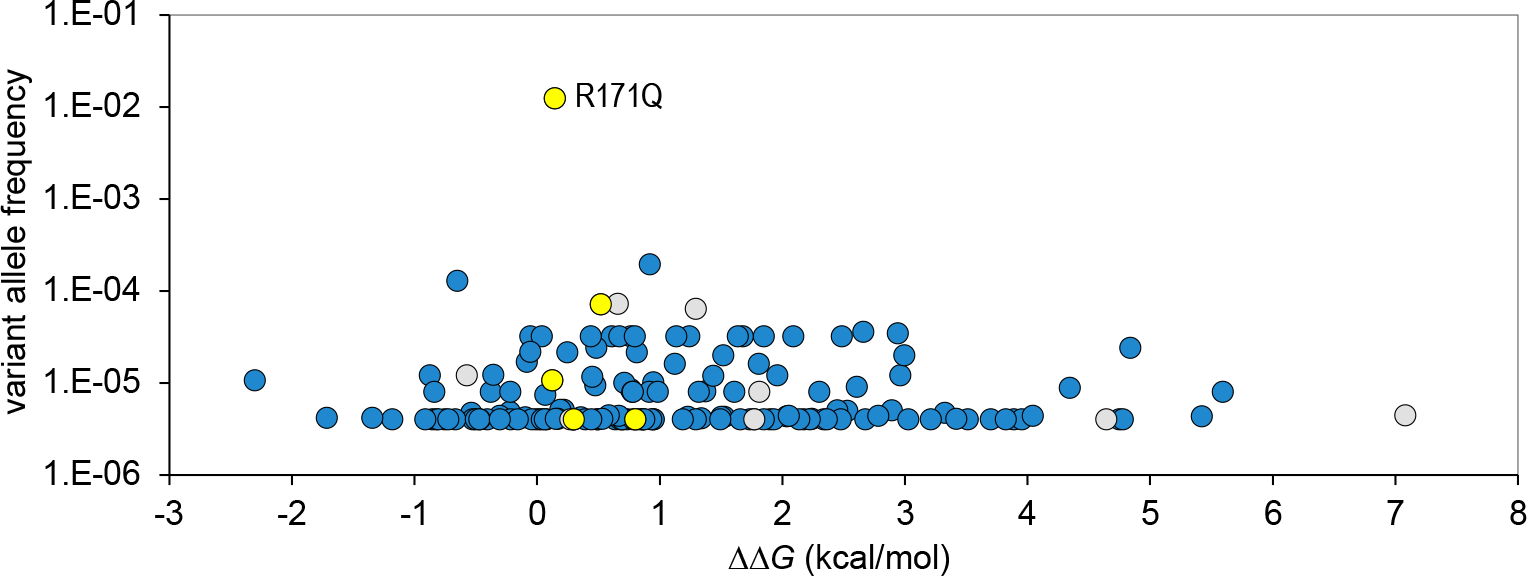
Population frequency of *MEN1* variants. The frequency of benign and uncertain missense variants in the gnomAD database plotted against ΔΔ*G* value; blue fill: variants occurring in the gnomAD database only; yellow fill: variants reported in the gnomAD and SGCD databases; grey fill: variants in both gnomAD and HGMD Pro (DM class) databases. In cases where different nucleotide substitutions give rise to the same amino acid change, frequency is shown as a total for all variant alleles.

### Most pathogenic variants are buried in the Menin structure

To examine whether there are differences in the spatial distribution of benign and pathogenic variants, we calculated the relative solvent accessibility (RSA) of wild-type residues at all positions of missense substitutions (Table 3). This showed that while positions of benign variants are distributed throughout the volume of the protein, 86.3% of pathogenic variants occur in solvent-inaccessible (i.e. buried) regions of RSA <0.2 (Figure 5A). Notably, this is also true for the 7 novel variants, 6 of which had an RSA value <0.02. Plotting RSA against ΔΔ*G* showed that variants at buried positions were also likely to be the most strongly destabilizing to protein structure (Figure 5B). Nevertheless, we observed that a significant number of pathogenic variants exhibited both accessibility to solvent (RSA>0.2) and relatively low ΔΔ*G*. Mapping the positions of solvent-accessible variants onto the surface of Menin showed that, as for distribution throughout the internal volume of the protein, benign variants tended to be distributed across the surface. In contrast, pathogenic variants appeared to occur in clusters, one of which corresponds to binding surfaces for JunD, KMT2A and PSIP (Figure 5C, D), while another occurs on the opposite surface of Menin to the JunD binding pocket. It is possible therefore that the latter region represents the site of an as-yet uncharacterized functional interaction of Menin. As described above, 6/7 novel missense variants occur at positions which are buried in the interior of the protein, whereas the only solvent-accessible variant, H46A, occurs at the interface with KMT2A and presumably acts to impair this interaction (Figure 5E).

**Figure 5:**
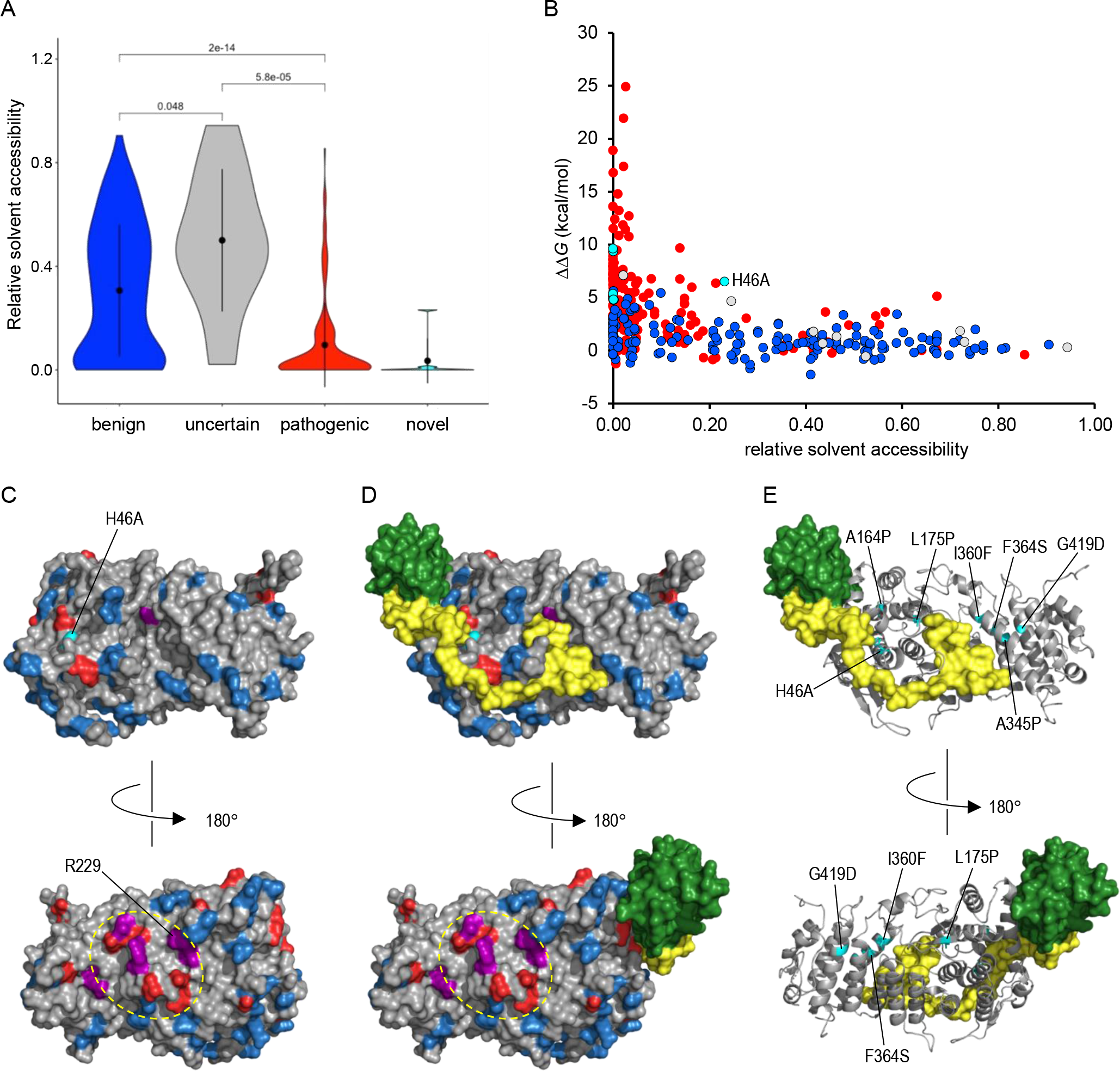
Molecular distribution of pathogenic and benign variants. A) Relative solvent accessibility was calculated for each variant group; black circles and vertical lines within each data area represent median and upper and lower quartiles respectively. Numbering above data points shows *p* values (Student’s Two-tailed t-test) between groups as indicated. B) Buried pathogenic variants are predicted to be the most destabilising to Menin structure; note that 6/7 of the novel missense variants reported here are deeply buried within the protein (RSA < 0.02), while only novel variant H46A is solvent accessible. C, D) Surface distribution of solvent-accessible variants. The surface of Menin (grey), either alone (C) or in complex (D) with KMT2A (yellow) and PSIP (green) shows all variants with RSA>0.2: blue, benign; red, pathogenic; purple colouring show positions at which different pathogenic and benign variants have been observed; the novel H46A variant is coloured cyan. The broken yellow oval indicates a cluster of pathogenic variants which may constitute an as yet unidentified interface for protein-protein interactions. E) Menin is shown as a grey ribbon; novel missense variants are coloured cyan with sidechains displayed in stick format; KMT2A and DSIP are shown as in D.

To investigate the effects of protein interactions on the thermodynamic effects of *MEN1* variants further, we compared ΔΔ*G* values for variants in PDB structure 3u88 (Menin complexed with KMT2A and PSIP peptides) by analysis both of Menin chains in isolation (chains A, B) and complexed to KMT2A and PSIP. As expected, regions of decreased solvent accessibility in the complexes aligned with residues annotated as forming protein-protein contacts (Figure 6). However, the presence of bound peptides had little effect on ΔΔ*G* values of benign variants, indicating that these have a neutral effect on protein binding. Conversely, protein binding had a large effect on ΔΔ*G* values of a number of pathogenic variants; again, these predominantly occurred at or close to protein interfaces, indicating that these variants are likely to have a direct effect on ligand binding by Menin.

**Figure 6:**
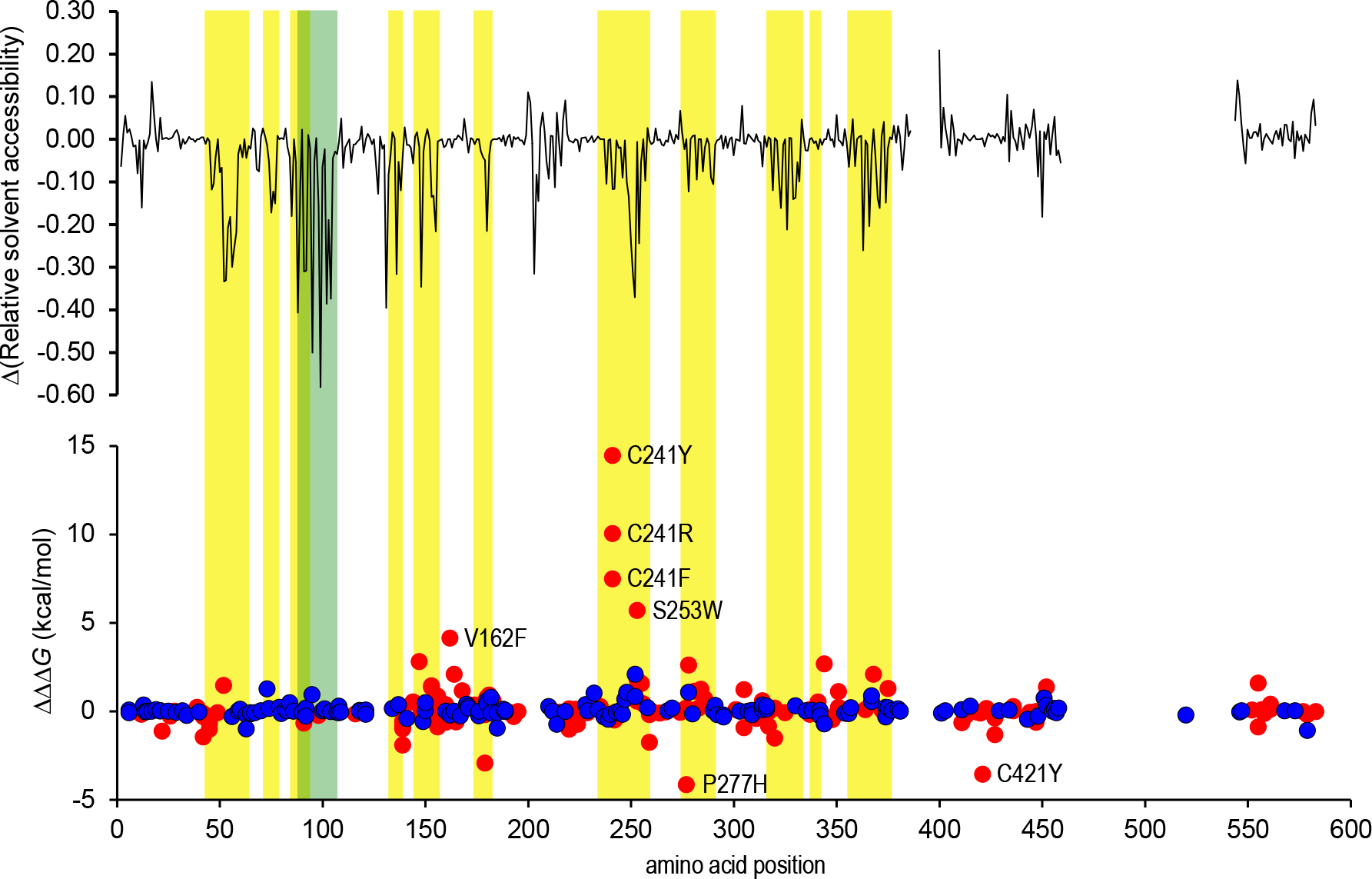
Effect of protein-protein interaction on ΔΔ*G*. Analysis of solvent accessibility and thermodynamic effect of variants was performed on PDB 3u88 (Menin:KMT2A:DSIP complex), both on Menin chains in isolation (chains A, B) and as part of the complex. The upper graph shows the average difference in solvent accessibility by position in the complexed and isolated Menin chains respectively (ΔRSA = RSA [complex] – RSA [isolated]); the lower graph shows the equivalent difference in average ΔΔ*G* value at each position (i.e. ΔΔΔ*G*); data points are labelled for variants where ΔΔΔ*G* 3 kcal/mol; background shading indicates positions of Menin residues forming contacts with KMT2A (yellow) or DSIP (green) in PDB 3u88.

### Destabilizing variants reduce levels of functional Menin protein

Previous reports studying the effects of missense variants on levels of functional Menin within the cell have shown that pathogenic variants have a tendency to increase protein turnover and/or reduce the steady-state level of protein, while benign variants tend to have no such effect (40, 41). We correlated the previously-reported effects of variants on levels of steady-state protein with average ΔΔ*G* values, and observed that variants which were predicted to be strongly destabilizing *in silico* (ΔΔ*G* >3 kcal/mol) exhibited significantly lower levels of steady-state protein in cell-based assays (p=0.0001, Figure 7), consistent with the hypothesis that variants with high ΔΔ*G* values reduce the biological activity of Menin.

**Figure 7.**
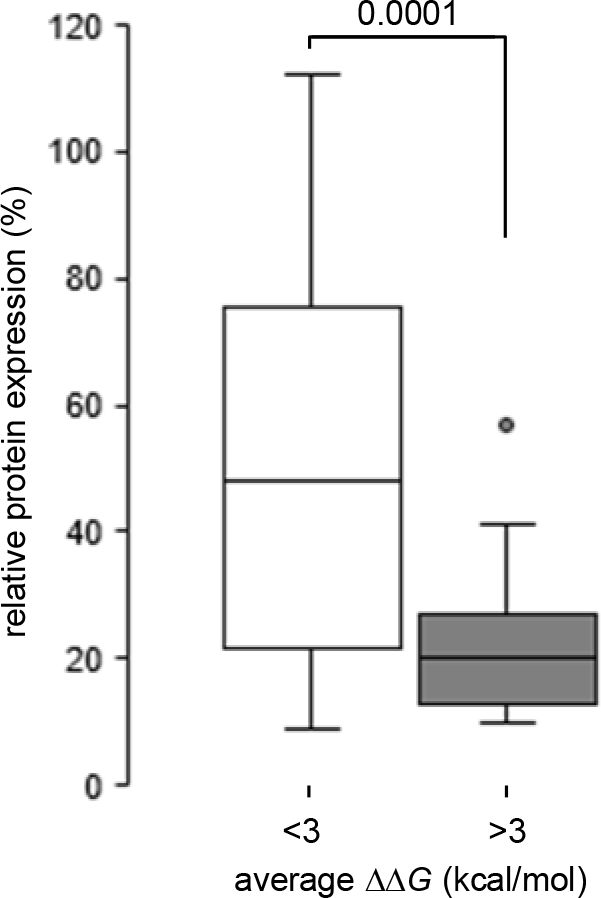
Predicted thermodynamic stability correlates with observed expression. A) Steady-state expression levels have been reported for a number of Menin variants; relative expression level data was sorted into two groups according to ΔΔ*G* value as calculated in this study (neutral or weakly destabilising: ΔΔ*G* <3 kcal/mol [n=14]; strongly destabilising: >3 kcal/mol [n=27]); boxes show the range between upper and lower quartiles; horizontal lines within each data box show median value; data points are shown for outliers only. The difference in relative expression between the two groups was highly significant (*p*=0.0001).

### Can ΔΔG value be used as an aid to variant classification?

To evaluate the clinical validity of ΔΔ*G* values, we performed Receiver Operating Characteristic (ROC) curve analysis for the groups of benign and pathogenic variants and compared the results with the outputs from a number of commonly-used phenotypic predictions tools: SIFT (42), PolyPhen (43) and REVEL (44). All methods yielded areas under the curve (AUC) of 0.819-0.864, indicating that all have clinical validity (Figure 8A). However ΔΔ*G* analysis resulted in the highest specificity but lowest sensitivity. Values of ΔΔ*G* > 3 kcal/mol are generally regarded as being strongly destabilizing towards protein structure (45); taking this as a threshold for variant classification gives sensitivity and specificity of 67.1% and 89.4% (positive predictive value, 86.4%), while setting a more conservative threshold of ≥4 kcal/mol yields increased the specificity to 95.0%, though with a concomitant loss of sensitivity (54.0%; positive predictive value, 90.6%). A marginal increase in positive predictive value (PPV) could be obtained by combining ΔΔ*G* thresholds with a cut-off in the REVEL score of 0.7, which has been reported to exclude 95% of false positive calls (46), yielding PPV’s of 87.7% at ΔΔ*G* ≥ 3 kcal/mol and 91.5% at ΔΔ*G* ≥ 4 kcal/mol. Notably, all seven novel missense variants reported here cluster within the upper right quadrant (Figure 8B), consistent with a severe impact on protein stability and suggesting that ΔΔ*G* values can potentially be used to provide evidence towards variant classification in MEN1.

**Figure 8.**
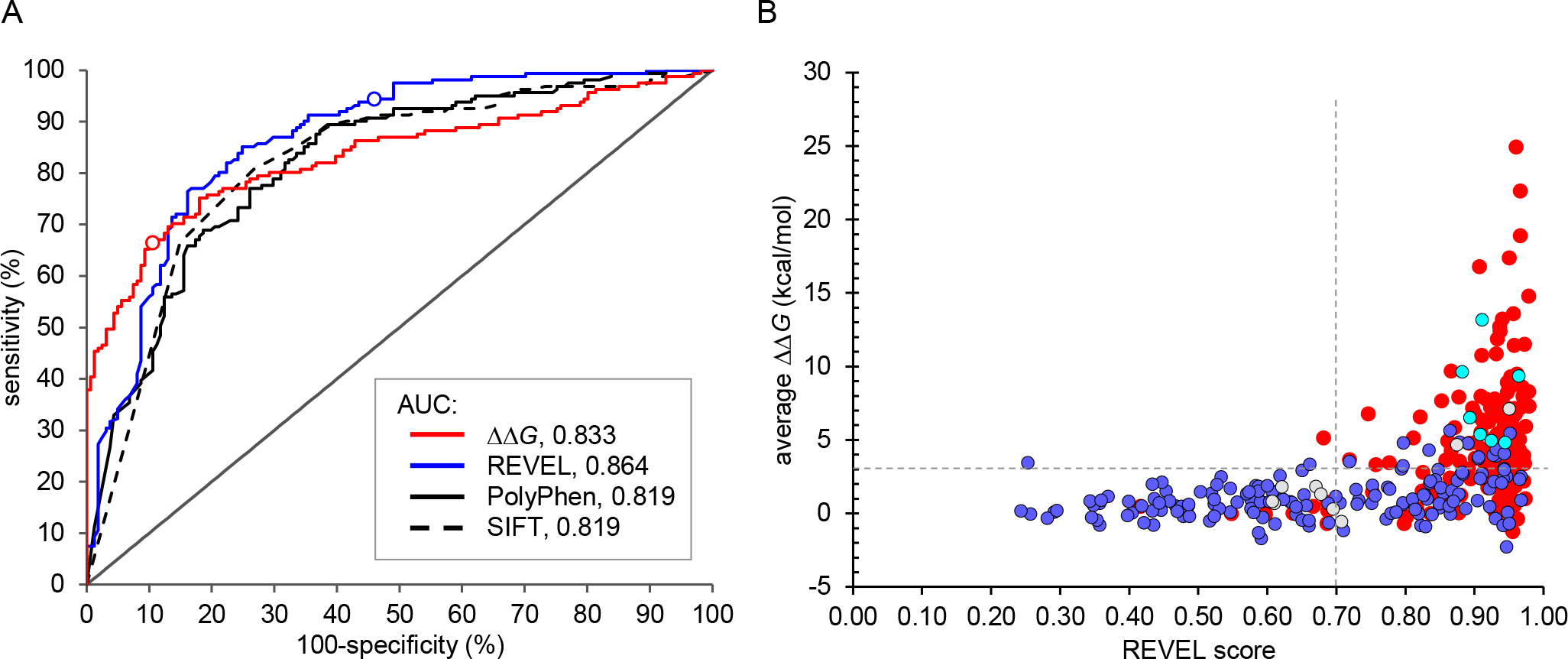
Using thermodynamic analysis to assess the impact of novel missense variants. A) ROC curves for groups of pathogenic and benign variants as functions of ΔΔ*G* value (red line; AUC, 0.833), REVEL score (blue line; AUC, 0.864) PolyPhen2 probability for pathogenicity (black line; AUC, 0.819) and SIFT score (broken black line; AUC = 0.819); open circles on ΔΔ*G* and REVEL traces indicate positions corresponding to threshold values of 3 kcal/mol and 0.7 respectively. B) Scatter plot of ΔΔ*G* value against REVEL score for all variants (red circles, pathogenic; blue fill, benign; grey fill, uncertain; cyan fill, novel). Where different nucleotide substitutions give rise to the same amino acid change, the REVEL score was calculated as an average of values for the individual nucleotide variants. Broken horizontal and vertical lines indicate thresholds of ΔΔ*G* = 3 kcal/mol and REVEL score = 0.7 respectively; note that all 7 novel missense variants cluster in the upper right quadrant of the plot.

## DISCUSSION

Previous work has shown that predicted thermodynamic destabilization of protein structure, as measured by ΔΔ*G* values calculated by FoldX, can be used as a predictor of pathogenicity in *MSH2* and *PAH* variants (7, 8). Our data indicates that the same is true for variants in *MEN1*, and that a high predicted ΔΔ*G* value is a strong positive predictor for pathogenicity. Using a threshold of only 3 kcal/mol, specificity for variant classification was 89.4%, rising to 95.0% for a more conservative threshold of 4 kcal/mol. By contrast, using a proposed threshold of 0.7 for the phenotypic meta-prediction tool REVEL yielded a specificity of only 53%. Since MEN1 has variable penetrance and often late onset, the identification of likely pathogenic variants has significant implications for patient surveillance and genetic testing of family members. With respect to the seven novel missense variants reported here, all had high average predicted ΔΔ*G* values (range, 4.81-13.16 kcal/mol) and six were deeply buried within the protein, strongly supporting pathogenicity. All these cases were also predicted as deleterious or probably pathogenic by commonly-used tools for *in silico* pathogenicity prediction; however, the comparatively low specificity of all these tools for variants in *MEN1* highlights the value of thermodynamic analysis as a means of reducing false positive calls.

As might be expected, our analysis shows that variants which are buried within the Menin structure are those that are predicted to result in greatest structural destabilization. In fact, the majority of reported pathogenic variants in *MEN1* are buried, suggesting that any novel variant which is solvent inaccessible (RSA<0.2) and has a predicted ΔΔ*G* >4 kcal/mol is also highly likely to be pathogenic.

Nevertheless, a number of pathogenic variants lie on the surface of Menin, and many of these have relatively low ΔΔ*G* values. A number of these variants lie at or close to positions of known interactions with binding partners such as JunD, KMT2A or PSIP, where they presumably have an adverse effect on binding of these factors, emphasizing the value of integrating all known structural annotation into a final classification of the likely effect of a variant. Our data also suggests the possible existence of an as-yet unidentified interaction of Menin, as evidenced by the cluster of pathogenic variants lying on the protein surface opposite the JunD binding pocket. Notably, *MEN1* has recently been identified as one of the genes exhibiting significant spatial clustering of pathogenic variants (47); our analysis suggests that this clustering is likely to apply both to regions of structural importance, which are buried in the interior of the protein, and to surface regions which form essential interactions with binding partners.

In terms of broader applicability of this approach, our work builds upon the reported analysis of *MSH2* and *PAH* variants and applies it to the classification of novel clinical variants. Whether the same approach can be used for other proteins remains to be determined. One obvious limitation of structural analysis is, by definition, the need for a suitable structural model. However, even where no experimental structures are available for a protein of interest, it may still be possible to use comparative modelling to generate a reliable model of regions or domains which can be used for structural analysis. Another likely limitation is the architecture of the protein itself. Both Menin and MSH2 are relatively compact, globular proteins, with low surface area to volume ratio and a high proportion of amino acids in regions of secondary structure. As a result, the effect of missense variants on the internal geometry and thermodynamic stability of the proteins is amenable to *in silico* prediction, particularly given the availability of suitable high-quality PDB structures. However, less well-structured proteins, or fibrillar proteins where a greater proportion of amino acids are exposed to solvent, are likely to be less amenable to such study as the confidence with which the structural and thermodynamic effects of missense variants can be predicted will be greatly reduced.

Such rules are likely to be revealed only by proteome-wide study which is beyond the scope of this manuscript.

In summary, we have shown that structural analysis of missense substitutions in *MEN1* can be used to identify variants likely to destabilize the protein and thus potentially as an aid in variant classification. Given that all analysis described herein used publicly-available data, freely-available software and does not require specialist bioinformatic skills or infrastructure, such analysis lies within the capability of any genetics laboratory or testing service. As such, there is significant scope to make greater use of protein structural data in the routine interpretation of genetic variation.

## ACKNOWLEDGMENTS

The authors wish to acknowledge support from the Wellcome Trust (grant no. 200990).

## DATA AVAILABILITY

All data generated or analyzed during this study are included in this published article, with the exception of ΔΔ*G* and RSA values shown in Table 3, which shows average values for each variant calculated from all PDB structures used in the analysis as indicated in the table. Full data are available from the corresponding author on reasonable request.

